# CDK9 pharmacological inhibition with PRT2527 has antitumor activity in marginal zone lymphoma models and can improve the effects of BTK, PI3K, and BCL2 inhibitors

**DOI:** 10.64898/2026.02.10.705084

**Authors:** Eleonora Cannas, Alberto J. Arribas, Luciano Cascione, Francesca Guidetti, Andrea Rinaldi, Davide Rossi, Anastasios Stathis, Diane Heiser, Francesco Bertoni

**Affiliations:** Institute of Oncology Research, Faculty of Biomedical Sciences, USI, Bellinzona, Switzerland; SIB Swiss Institute of Bioinformatics, Lausanne, Switzerland; Oncology Institute of Southern Switzerland, Ente Ospedaliero Cantonale, Bellinzona, Switzerland; Prelude Therapeutics Inc, Wilmington, DE, USA

## Abstract

Cyclin-dependent kinase 9 (CDK9) drives transcriptional elongation and supports the expression of short-lived oncogenic and anti-apoptotic proteins such as MYC and MCL1. PRT2527 is a potent, selective CDK9 inhibitor currently in early clinical development. We evaluated its preclinical activity in marginal zone lymphoma (MZL) models, including cell lines with acquired resistance to BTK, PI3K, and BCL2 inhibitors. Short exposure (4 hours) to PRT2527 produced nanomolar cytotoxicity across all tested MZL cell lines, with efficacy maintained in resistant derivatives. Transcriptomic profiling of VL51 cells showed broad gene repression, including MYC, IRF4, NF-κB-related genes, and MCL1, alongside increased expression of HLA class II genes. Moreover, comparison with additional CDK inhibitors revealed a similar transcriptional repression signature, underscoring a conserved CDK-dependent regulatory network. Protein analyses confirmed rapid depletion of MCL1, MYC, RNA polymerase II, and IRF4. Flow cytometry validated increased HLA class II and decreased HLA class I surface expression. Combination studies demonstrated additive to synergistic effects with BTK inhibition (ibrutinib) or dual PI3K/BCL2 inhibition (copanlisib plus venetoclax), independent of baseline drug sensitivity. Mechanistically, these combinations may enhance apoptosis by concurrently suppressing survival signaling and transcriptional addiction. Analysis of patient samples revealed high CDK9 expression, further supporting the biological relevance and therapeutic rationale for targeting CDK9 in this disease. Our findings support the development of CDK9-based combination strategies for relapsed/refractory MZL and other B-cell malignancies, with an additional potential for integration with immunotherapies.

**Key Points:**

1. CDK9 inhibitor PRT2527 kills marginal zone lymphoma cells, regardless of whether they are resistant to other targeted drugs.
2. PRT2527 boosts the effects of BTK, PI3K, and BCL2 inhibitors and alters immune-related gene expression.

**Draft of graphical abstract:** 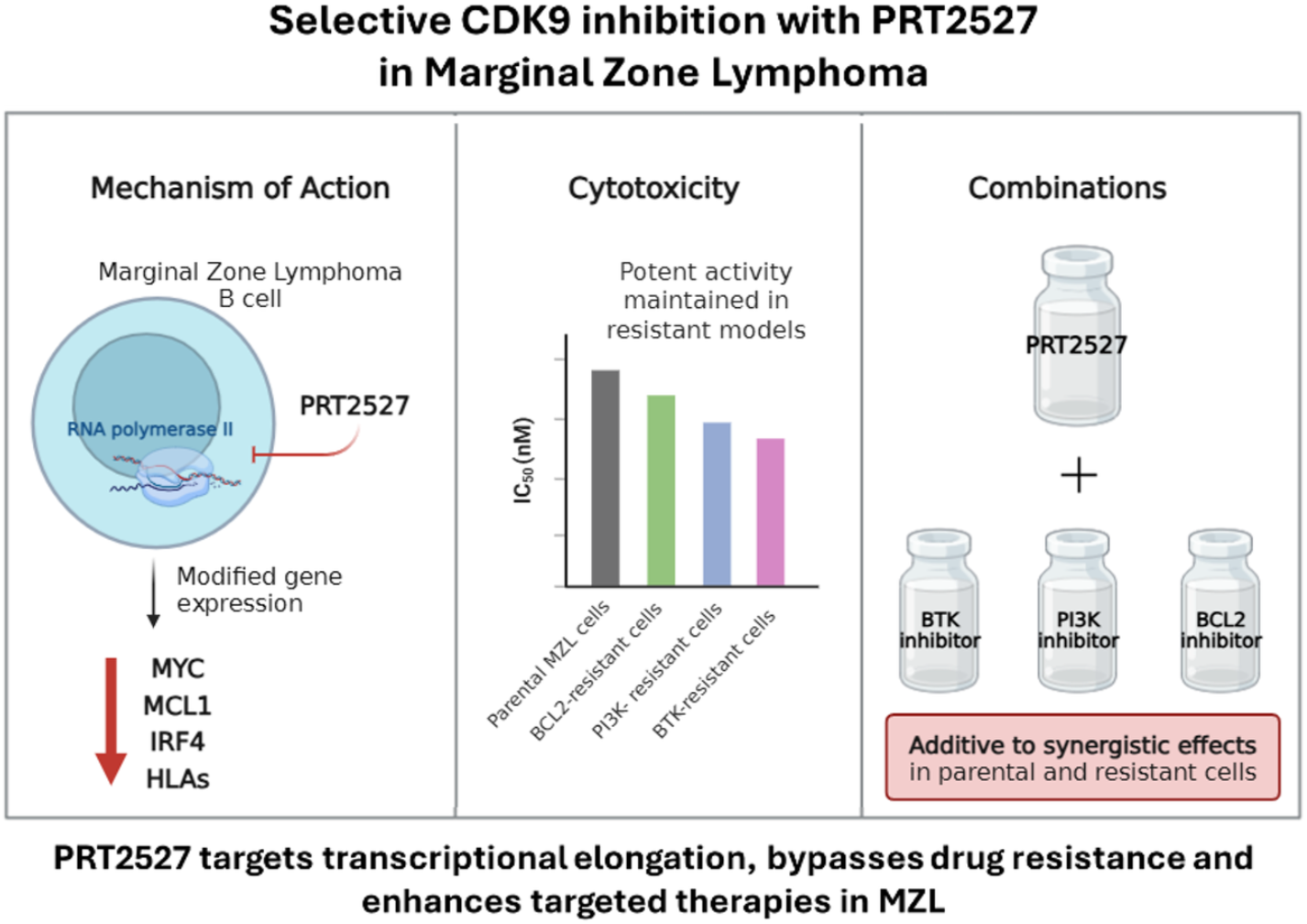

## Introduction

Cyclin-dependent kinase 9 (CDK9) is often abnormally expressed in both solid tumors and hematological cancers ^1-4^. This is not surprising, as CDK9 is the catalytic core of positive transcription elongation factor b (P-TEFb), which primarily partners with cyclin T1/T2 to phosphorylate the RNA polymerase II C-terminal domain (Ser2) and to release paused polymerase into productive elongation by phosphorylating negative elongation factors NELF/DSIF (SPT5) and related cofactors ^1-3,5-7^. This pause-release step functions as a rate-limiting checkpoint for rapid gene expression programs, supporting the integrity of superenhancer-driven oncogenic transcription. Consequently, acute inhibition of CDK9 quickly causes the collapse of short-lived mRNAs and proteins, particularly MYC and the anti-apoptotic BCL2 family members MCL1 and BFL1, shifting the balance toward apoptosis in transcription-dependent cancers. Additionally, CDK9 plays a role in epigenetic regulation by phosphorylating the helicase Brahma-related gene 1 (BRG1), a key component of the SWI/SNF complex ^2^, and is involved in the DNA damage response through its interaction with cyclin K ^8^.

In normal lymphocytes, CDK9 is expressed at various stages of development, including pro-B cells, a small fraction of pre-B cells, germinal center B cells (particularly centroblasts), and some scattered B-cell blasts in the interfollicular areas ^4^. Indeed, B cells are especially dependent on rapid transcriptional reprogramming during germinal center reactions and plasma cell differentiation; here, MYC pulses and NF-κB/IRF-driven programs require efficient elongation ^9,10^. While physiological CDK9 activity is critical for these transitions, it is not required for quiescent states, creating an exploitable therapeutic window when malignant B cells become transcriptionally dependent. We and others have shown that targeting cell-cycle- and pan-CDK inhibitors results in preclinical anti-lymphoma activity ^11-15^. Mechanistically, in B-cell malignancies, CDK9 inhibition enforces promoter-proximal pausing, rapidly depletes MYC and MCL1, and induces mitochondrial apoptosis ^12,13,15,16^. These effects have been observed across diffuse large B-cell lymphoma (DLBCL), mantle cell lymphoma (MCL), and chronic lymphocytic leukemia (CLL) models and primary cells ^16-19^.

Historically, pan-CDK agents showed activity but had limited clinical tolerability due to off-target myelosuppression and gastrointestinal toxicities ^17,18,20^. Newer-generation, CDK9-selective inhibitors have been designed to widen the therapeutic index while preserving effects on short-half-life survival signals ^3,17,18,20,21^. CDK9-selective inhibitors, such as istisociclib (KB-0742) ^22^, fadraciclib (CYC065)^23^, voruciclib (ME-522)^24^, enitociclib (BAY-1251152; VIP-152) ^25^, tambiciclib (SLS009/GFH009) ^26^, QHRD107 ^27^, and PRT2527 ^28,29^, have entered early clinical development and appear well tolerated, with manageable toxicity. Among these, PRT2527 is a highly potent, ATP-competitive, and kinase-selective CDK9 inhibitor ^30^. Preclinically, PRT2527 reduces MYC and MCL1 protein levels and shows robust activity in DLBCL, MCL, and CLL models. Combinations with BTK or BCL2 inhibitors deepen and prolong responses, consistent with convergent control of BCR-survival signaling and mitochondrial priming ^31^. Clinical activity and an acceptable safety profile as monotherapy and in combination with zanubrutinib have been reported in the first 56 patients enrolled in the phase 1 study for relapsed/refractory lymphoma patients ^28^. The most common treatment-related adverse event was neutropenia, which was managed with growth factor support ^28^.

This manuscript presents a comprehensive preclinical and translational evaluation of PRT2527 as a single agent and in combination with other agents in marginal zone lymphoma (MZL) models, including cell lines with acquired resistance to BTK, BCL2, and PI3K inhibitors.

## Materials and methods

### Cell lines

Cell lines (VL51 ^32^, SSK41 ^33^, Karpas1718 ^32^, HC1, ESKOL, HAIR-M ^34^, VL51-Ide ^35^, VL51-Ibru ^36^, VL51-Copa ^37^, SSK41-CV14 ^38^, Karpas1718-Ide ^39^) were cultured under recommended conditions. All media were supplemented with fetal bovine serum (FBS) (10% or 20%) and penicillin-streptomycin-neomycin (≈5,000 units penicillin, 5 mg streptomycin, and 10 mg neomycin/mL; Sigma-Aldrich, Darmstadt, Germany). Cell line identities were confirmed by short tandem repeat DNA fingerprinting using the Promega GenePrint 10 System kit (B9510). Cells were periodically tested for mycoplasma negativity using the MycoAlert Mycoplasma Detection Kit (Lonza, Visp, Switzerland).

### In vitro cytotoxic activity

Cells were seeded at 10,000 or 20,000 cells per well in 96-well plates and exposed to PRT2527 in a 1:3 dilution series from 0.01 μM to 10 μM for 4 hours. After compound washout, cells were assayed by MTT [3-(4,5-dimethylthiazolyl-2)-2,5-diphenyltetrazoliumbromide] two days after initial treatment. PRT2527 was combined with venetoclax, copanlisib, idelalisib, and ibrutinib for two or three days, followed by an MTT assay. PRT2527 was kindly provided by Prelude. Venetoclax, copanlisib, idelalisib, and ibrutinib were purchased from Selleckchem (Houston, TX, USA). Drug doses and dilutions are shown in Supplementary Table S1.

Sensitivity to drug treatments was assessed using IC50 values, calculated via 4-parameter logistic regression in the PharmacoGx R package ^40^. The beneficial effect of the combinations relative to single agents was evaluated using both the Chou-Talalay combination index ^41^, and the Highest Single Agent (HSA) model as calculated with the SynergyFinder R package ^42^.

### Transcriptome analysis and data mining

Cells were seeded at 500,000 cells per well in a 6-well plate and exposed to the compound or DMSO. RNA was extracted and processed for RNA sequencing (RNA-Seq; stranded, single-ended 120 bp reads) using the NEBNext Ultra Directional RNA Library Prep Kit for Illumina (New England BioLabs Inc., Ipswich, MA, USA) on a NextSeq 2000 (Illumina, San Diego, CA, USA). Read quality was evaluated with FastQC (v0.11.5), and low-quality reads and bases, along with adaptor sequences, were removed using Trimmomatic (v0.35). High-quality reads were trimmed and aligned to the reference genome (HG38) using STAR. Gene expression was quantified using the HTSeq-count software package, with GENCODE v22 as the gene annotation. Expression values are provided in a tab-delimited format. Data were subset to genes with a counts-per-million value greater than five in at least one sample. The data were normalized using the ‘TMM’ method from the edgeR package and transformed to log2 counts per million using the edgeR function ‘cpm’. Differential gene expression for each comparison of interest was computed using a moderated t-test based on TMM normalization and voom transformation. The false discovery rate (FDR, as calculated using the Benjamin-Hochberg correction) was used to control for false positives. FDR <0.001 and an absolute fold-change greater than 1 (RNA-seq). Functional analysis was performed using GSEA (Gene Set Enrichment Analysis) with the MSigDB (Molecular Signatures Database) C2-C7 gene sets ^43^ and the SignatureDB database (https://lymphochip.nih.gov/signaturedb/). Statistical tests were performed using the R environment (R Studio console; RStudio, Boston, MA, USA). Unsupervised multidimensional scaling plots were used to visualize the parental multi-omics profile.

RNA-Seq data are available at the National Center for Biotechnology Information (NCBI) Gene Expression Omnibus (GEO; http://www.ncbi.nlm.nih.gov/geo/) under accession number GSE313143.

### HLA expression

Cells were seeded at 250,000 cells per well in a 6-well plate and then treated with varying concentrations of the compound for 24 hours. Cells were pre-incubated with Human TruStain FcX (Fc Receptor Blocking Solution, BioLegend, cat. No. 422301) for 10 minutes and stained with APC anti-human HLA A, B, C (BioLegend, cat. 311410), FITC anti-human HLA DR, DP, DQ (BioLegend, cat. 307632), or their respective mouse IgG2a isotype controls (BioLegend, APC cat. 400220, FITC cat. 400210). Acquisitions were performed on a FACSFortessa instrument (BD Biosciences, Allschwil, Switzerland), and the data were analyzed using FlowJo software (TreeStar Inc., Ashland, OR, USA).

### Immunoblotting

Cells were seeded in T25 flasks at a density of 5 million cells per 10 mL and treated for 24 hours with DMSO or varying concentrations of the compound. Subsequently, protein extraction was performed by incubating all conditions with M-PER (Mammalian Protein Extraction Reagent) lysis buffer plus Halt Protease and Phosphatase Inhibitor Cocktail, EDTA-free (100X), for 30 minutes on ice, followed by centrifugation at high speed at 4°C for 30 minutes. The protein concentration was determined using the BCA protein assay (Pierce Chemical Co., Dallas, TX, USA), and 50 µg of total protein was loaded per sample. Cell lysates were separated by SDS-PAGE on a 4-20% SDS-polyacrylamide gradient gel, and the proteins were transferred to nitrocellulose membranes for analysis. Membranes were incubated overnight with primary antibodies, followed by incubation with anti-mouse or anti-rabbit secondary antibodies for 1 hour at room temperature. Enhanced chemiluminescence detection was performed following the manufacturer’s instructions (Amersham Life Science, Buckinghamshire, UK). Luminescence was measured using the Fusion Solo S instrument (Witec AG, Sursee, Switzerland). Protein quantification was performed using the Fusion Solo S instrument (Witec AG). Equal loading of samples was confirmed by probing for vinculin as a housekeeping gene. The antibodies used for the experiment were: anti-MCL1 (D35A5, CST-5453; Cell Signaling), anti-c-MYC (D84C12, CST-5605; Cell Signaling), anti-IRF4 (D43H10, CST-4299; Cell Signaling), anti-RNA polymerase II (8WG16; MMS-126R-500; Covance), and anti-vinculin (E1E9V, CST-13901; Cell Signaling).

### CDK9 expression

Raw counts for gene expression data from a cohort of splenic MZL patients were obtained from previously published datasets ^44^. Counts were normalized to transcript per million (TPM) to compare selected genes. An additional public dataset ^45^ containing clinical samples from patients with follicular lymphoma (FL), extranodal MZL, nodal MZL, and splenic MZL was used to examine CDK9 gene expression, along with control genes expected to be expressed (CD19) or not expressed (MYH7) in lymphoma. DNA methylation profiling data from SMZL patients and non-tumoral spleen samples were also obtained from public datasets ^46^ to compare the β-values of CDK9 methylation between patient samples and healthy spleen controls.

## Results

### The CDK9 inhibitor PRT2527 has dose-dependent activity in MZL models, including cells with secondary resistance to targeted agents

The effect of the CDK9 inhibitor PRT2527 on cell viability was first assessed in six MZL-derived cell lines (Figure 1A). Based on the human pharmacokinetics of PRT2527 ^29^, the drug was applied for 4 hours, after which the cells were cultured in drug-free medium for 2 days before measuring the anti-tumor response. The compound showed dose-dependent activity across all tested cell lines, with IC50 values ranging from 186 nM to 1.3 μM (median, 580 nM). Notably, this activity persisted in derivatives with secondary resistance to BTK, BCL2, and PI3K inhibitors (Figure 1B).

**Figure 1.**
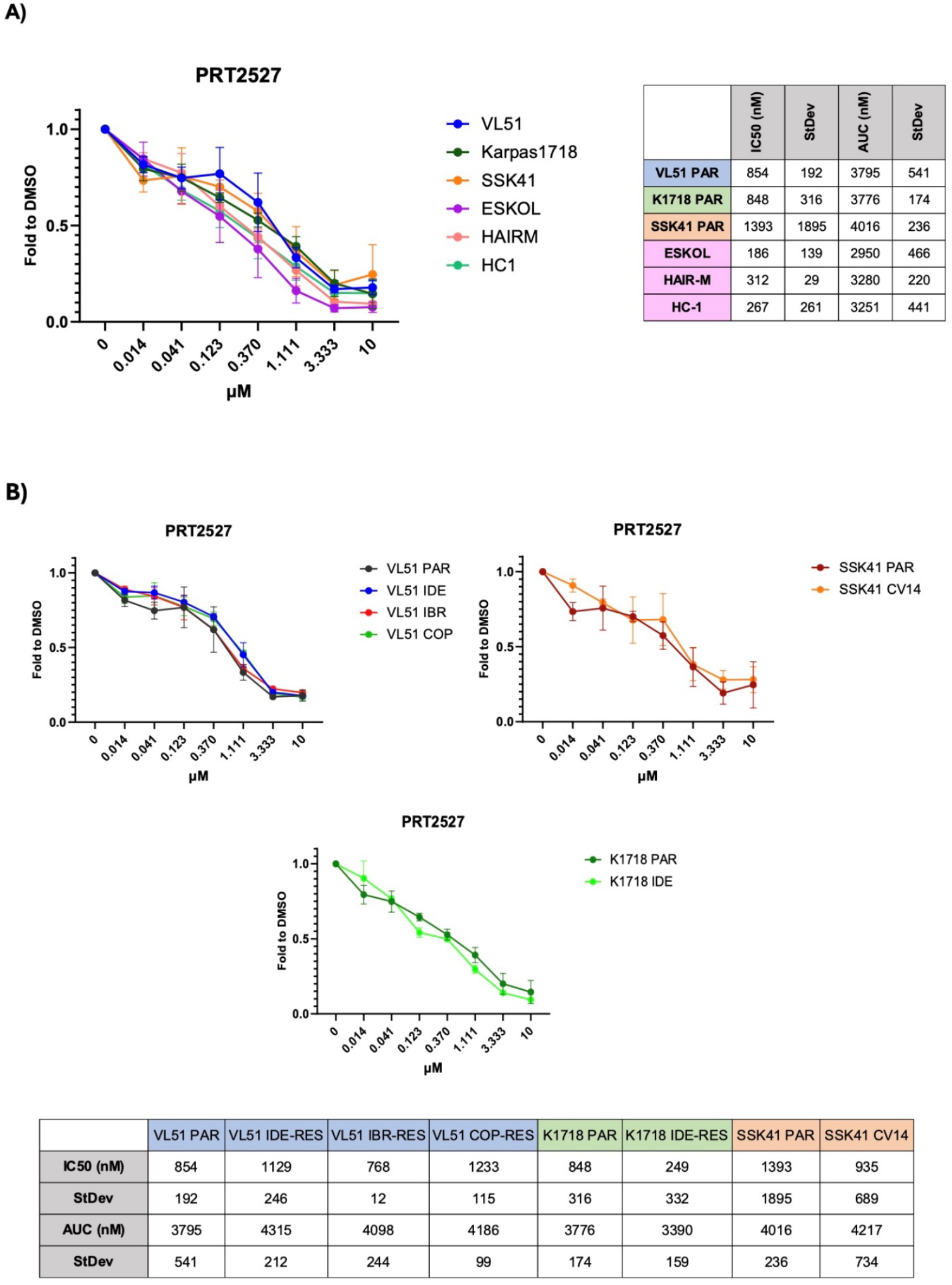
Effect of four hours of exposure to the CDK9 inhibitor PRT2527 in MZL models (A) and their derivatives with acquired resistance to PI3K, BTK, and BCL2 inhibitors (B), followed by MTT assay. MTT is read after two days following the initial drug exposure.

### The CDK9 inhibitor PRT2527 induces strong transcriptome changes in VL51 cells

To better understand the mechanism of action of the CDK9 inhibitor in our models, we analyzed the gene expression profile by RNA-seq in VL51 cells treated with PRT2527 at its IC50 (850 nM) or, as a control, with DMSO for 4 hours (Supplementary Table S2). As shown in Figure 2A, the transcriptomic landscape induced by CDK9 inhibition differed markedly from that of the control. Consistent with its role as an RNA polymerase II inhibitor, PRT2527 downregulated a substantially larger number of genes than it upregulated. Specifically, the CDK9 inhibitor repressed key genes involved in B cell proliferation and immune responses (*MYC, DUSP5, IRF2BP2, IRF4*, and *GPR183*), as well as anti-apoptotic pathways (NF-κB and MCL-1). In contrast, PRT2527 induced expression of genes primarily associated with glucose metabolism (*PDK1, P4HA1*, and *ENO2*). Among the upregulated transcripts were several HLA genes (HLA-DRB1, HLA-DRB5, and HLA-DPB1), which encode components of the major histocompatibility complex class II (MHC II) (Figure 2B). GSEA functional analysis revealed enrichment of gene sets related to cellular metabolism, MHC pathways, oxidative phosphorylation, and translation elongation. In contrast, gene sets associated with apoptosis, cell cycle progression, the NF-κB pathway, and MYC targets were significantly downregulated (Figure 2C). Notably, target genes of RNA polymerase II were strongly repressed, consistent with the drug’s mechanism of action. To further contextualize the transcriptional response, we used L1000 signatures in GSEA to identify compounds that generate similar or contrasting expression patterns relative to PRT2527 (Figure 2D). The positively enriched signatures included other CDK inhibitors (triptolide, AZD-5438, alvocidib, and SNS-032), as well as MAPK inhibitors (JNK-9L and JNK Inhibitor V) and MEK inhibitors (gomekli and AZD-8330). Conversely, PRT2527 showed a negative correlation with signatures associated with protein synthesis inhibitors (emetine and anisomycin), tubulin polymerization inhibitors (homoharringtonine), ALK inhibitors (LDN-193189), and ubiquitin protease inhibitors (NSC-632839).

**Figure 2.**
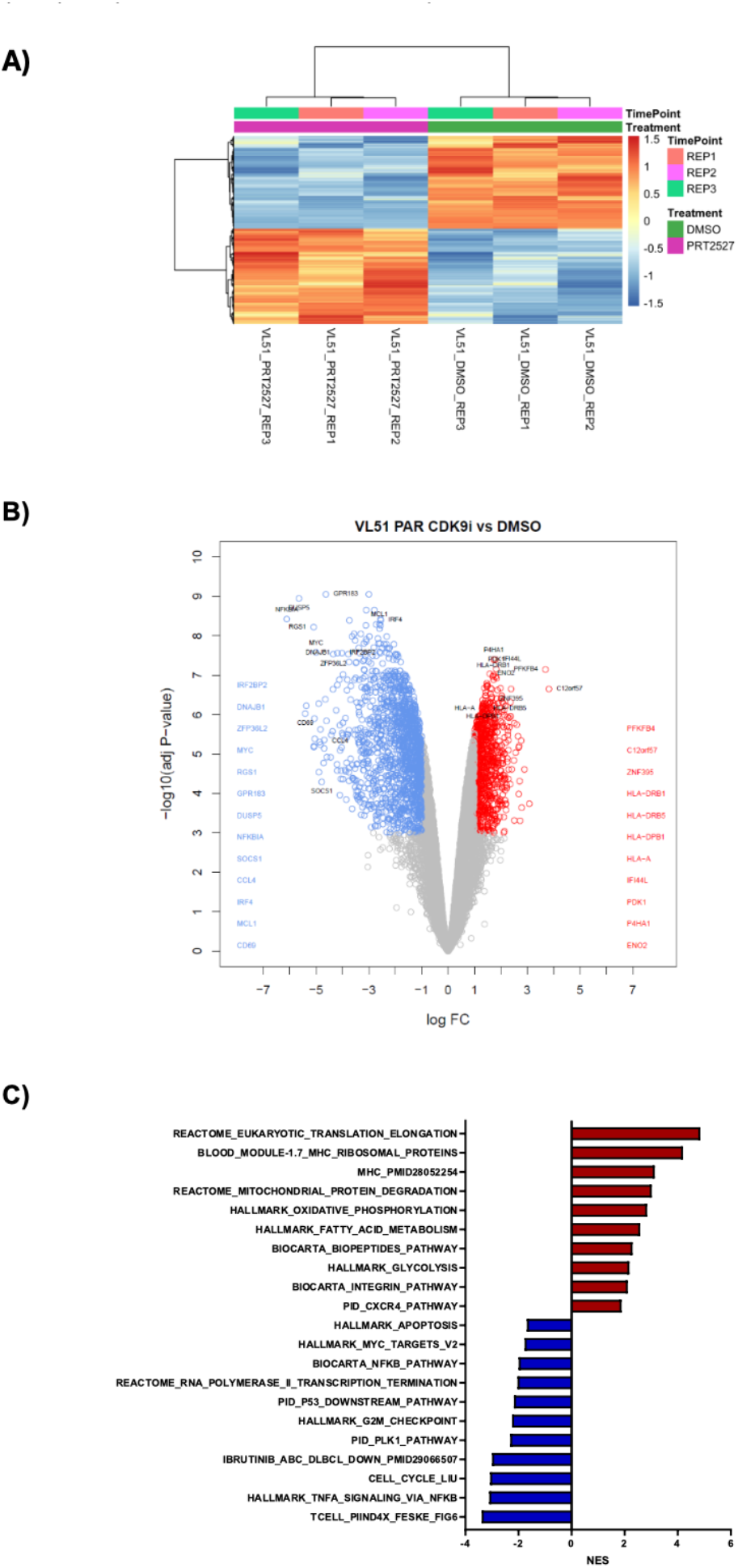
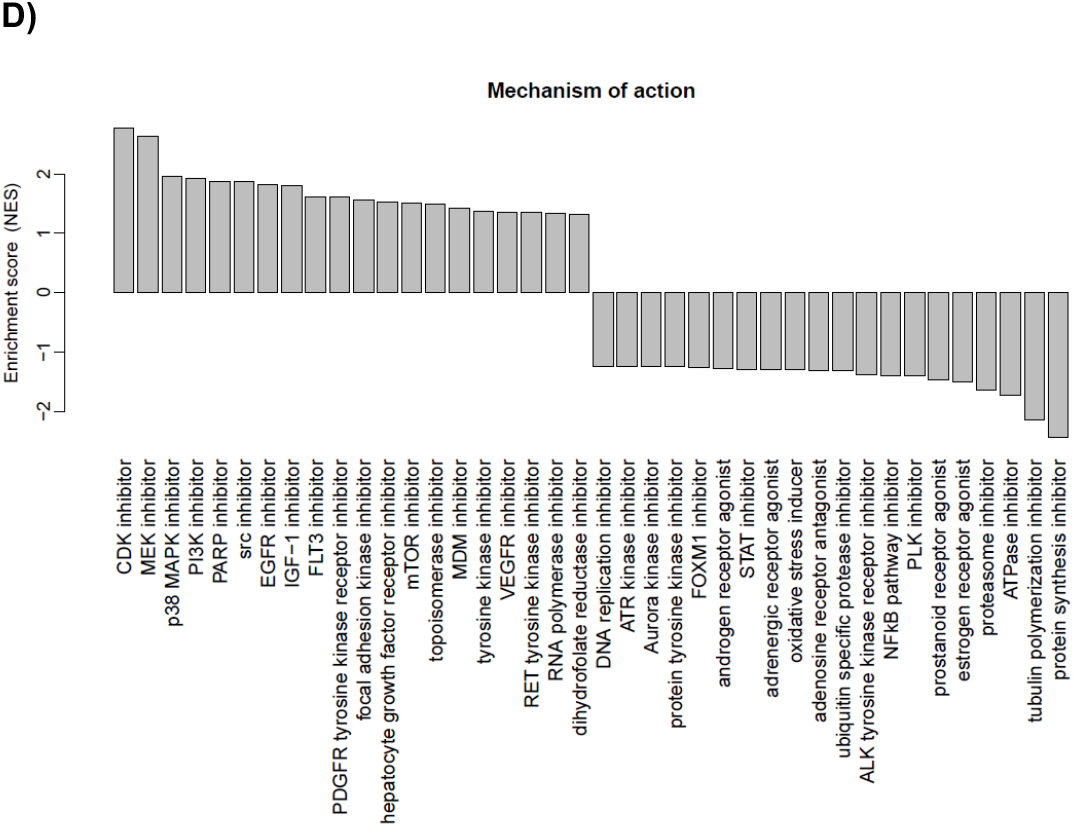
Gene expression profiles (RNA-seq) of PRT2527 in the MZL cell line VL51 exposed to PRT2527 (IC50, 850 nM) compared to DMSO (control). Cells were treated for 4 hours in three independent replicates. **A)** Heatmap of the transcriptome profiles of the three replicates of DMSO and the PRT2527. **B)** Vulcano-plot with the differentially expressed transcripts (protein coding transcripts only), highlighted genes upon |logFC|>1 and adj.P<0.001. Overexpressed genes are represented in red, and repressed genes are represented in blue. **C)** Gene Set Enrichment Analyses with the most differentially enriched gene sets (|NES|>1, P<0.05, and FDR<0.05). **D)** L1000 analyses with the drugs that give a similar or different pattern of expressed genes compared to PRT2527 (|NES|>1, P<0.05, and FDR<0.05).

### The CDK9 inhibitor PRT2527 downregulates the expression of pro-survival proteins in VL51 cells

We examined changes at the protein level of RNA polymerase II, c-MYC, IRF-4, and MCL-1 in the VL51 MZL model, exposing the cells to PRT2527 at 100 nM, 400 nM, and 850 nM. Low doses of the compound decreased c-MYC and eliminated MCL-1 levels (Figure 3). In fact, MCL-1 was entirely lost after exposure to the lowest concentration of the CDK9 inhibitor. Consistent with the transcriptome results, PRT2527 caused a strong reduction in RNA polymerase II levels and a significant decrease in IRF-4 at 100 nM, with no notable effects at higher doses.

**Figure 3.**
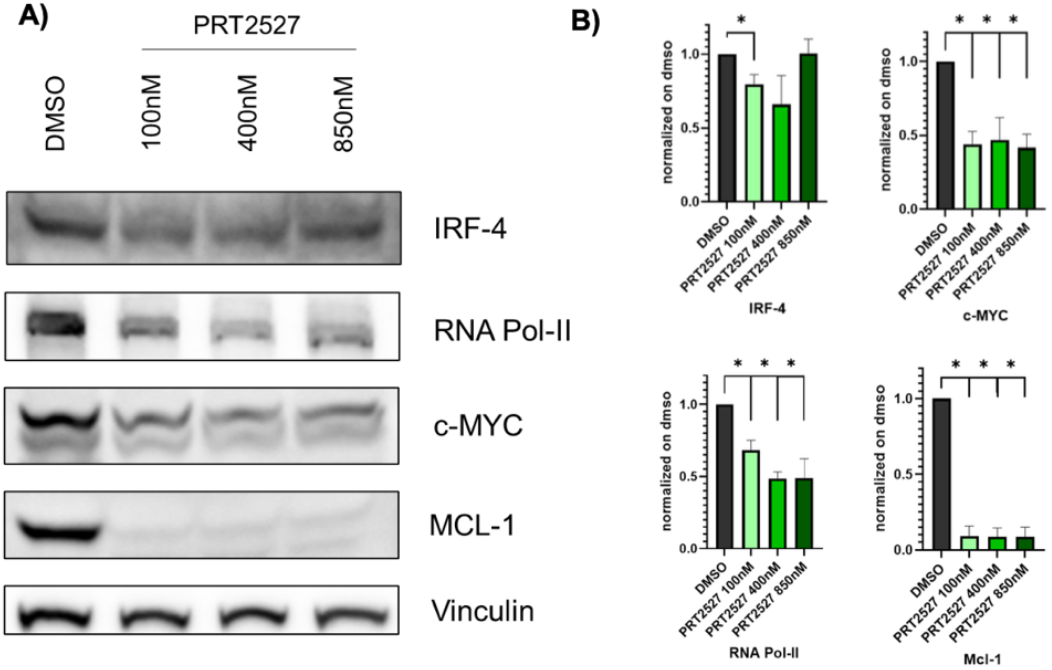
Immunoblotting of VL51 parental cells after exposure to PRT2527 at different concentrations (100nM, 400nM, or 850nM), or DMSO (control). **A)** Representative membranes of c-MYC, IRF-4, RNA Polymerase II, and MCL-1 protein expression after exposure to PRT2527 or DMSO (control) for 24 hours. **B)** Barplots represent the mean of protein quantification (normalized to vinculin) from at least two independent experiments, with error bars for standard deviation. * for p<0.05 in a Mann-Whitney U-test.

### The CDK9 inhibitor PRT2527 affects the surface expression of HLA class I molecules in MZL cell lines

To validate the RNA-Seq results, we evaluated surface expression of HLA class I (HLA-A, B, and C) and class II (HLA-DP, DR, and DQ) on VL51 cells after 24 hours of treatment with DMSO or increasing concentrations of the CDK9 inhibitor (100 nM, 400 nM, or 850 nM). As shown in Figure 4, HLA class II expression was significantly increased compared with the control at 100 nM of PRT2527 and remained elevated at higher doses. Conversely, HLA class I expression was significantly decreased by PRT2527 in a dose-dependent manner, consistent with the RNA-Seq gene expression profile (Supplementary Figure 1). To assess whether the effects of PRT2527 on HLA surface expression indicate a broader impact of CDK inhibition, we extended the analysis to additional MZL preclinical models (HAIR-M, ESKOL, and Karpas1718). In two of the three cell lines (ESKOL and Karpas1718), HLA class II expression increased, whereas in HAIR-M, we observed upregulation of HLA class I expression with no detectable changes in HLA class II surface expression (Supplementary Figure 2).

**Figure 4.**
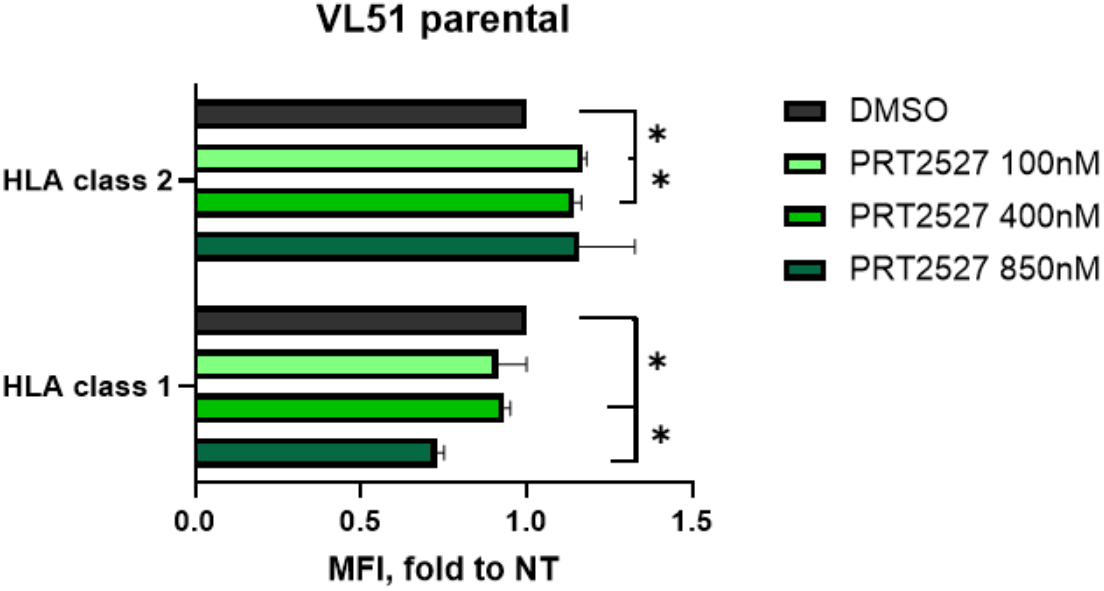
Surface HLA expression in VL51 parental cells after exposure to PRT2527 at different concentrations (100nM, 400nM, or 850nM) or DMSO (control). Cells were treated for 24 hours. Barplots represent the mean of at least two independent experiments, with error bars for standard deviation. * for p<0.05 in a Mann-Whitney U-test.

### The addition of PRT2527 increases the activity of BTK, PI3K, and BCL2 inhibitors

To explore the potential benefit of PRT2527-based combinations, we exposed three MZL cell lines and their respective resistant derivatives to increasing concentrations of the CDK9 inhibitor for 4 hours. After washout, we treated the cells with the BTK inhibitor ibrutinib, the PI3K inhibitor copanlisib plus the BCL2 inhibitor venetoclax, or the PI3K inhibitor idelalisib (Supplementary Table S1; Figure 5). Considering additivity and synergism, most combinations were beneficial across the parental cell lines, with limited benefit in the resistant lines. The best results were obtained when the CDK9 inhibitor was combined with ibrutinib or with copanlisib plus venetoclax. Based on this, we further explored the mechanism underlying the synergy of the combination of PRT2527 and ibrutinib or copanlisib/venetoclax for three days. In both cases, the combinations were beneficial, with no significant differences between the parental and their respective resistant derivatives. Overall, the effect of the two combinations was additive, with peaks of synergism between 10 nM and 100 nM of PRT2527 (Figure 5).

**Figure 5.**
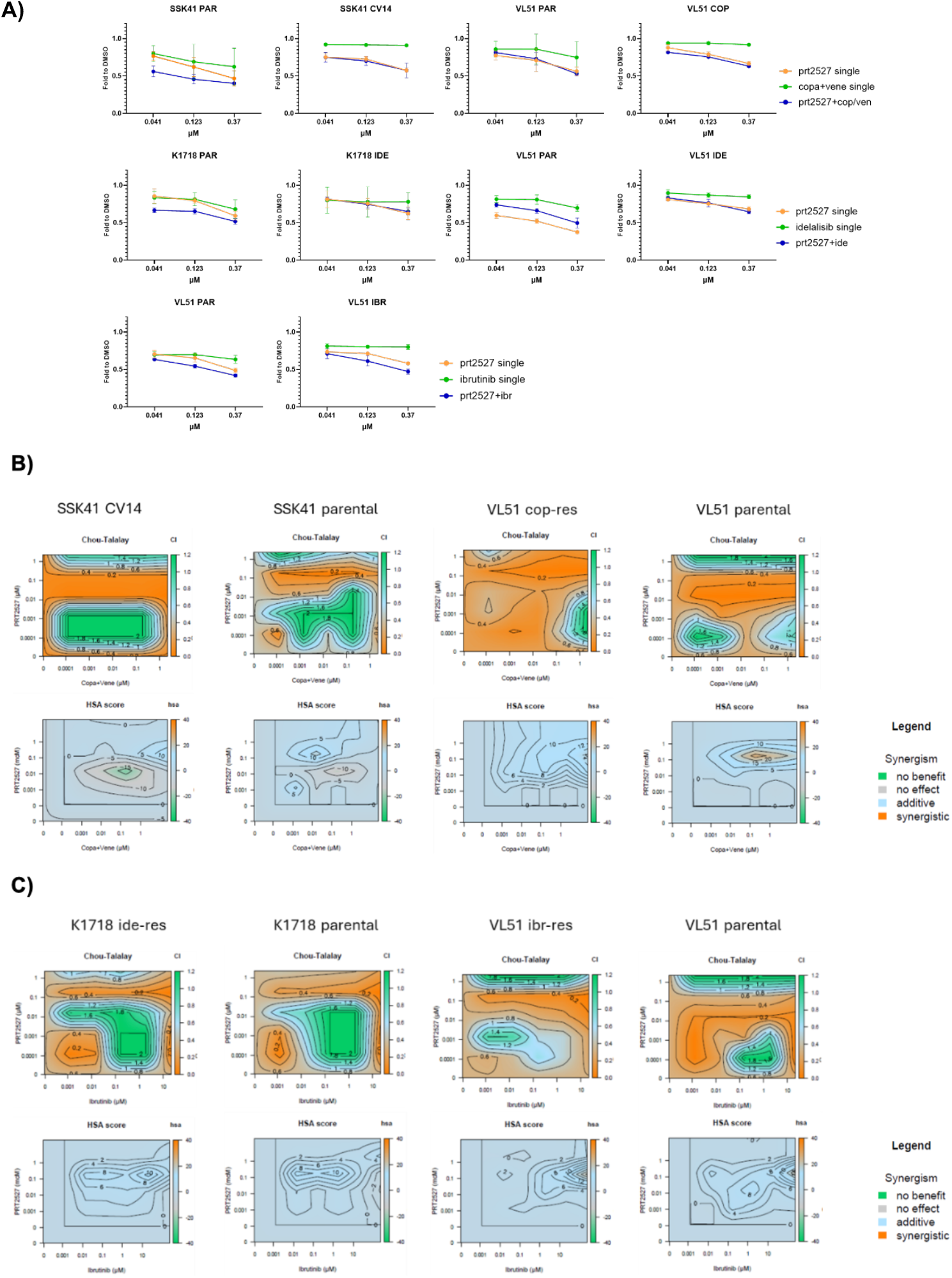
Effect of the CDK9 inhibitor PRT2527 in models with acquired resistance to PI3K, BTK, and PI3K/BCL2 inhibitors when combined with the drugs they are resistant to. **A)** Summary of the combinations of PRT2527 and copanlisib plus venetoclax, idelalisib, or ibrutinib, based on the drug to which the cell lines are resistant. Four hours of exposure to PRT2527, followed by washout and 48 hours of exposure to the second drug. MTT assay at the end of the treatment. **(B)** Heatmaps on the Chou-Talalay and HSA synergy models on the combination of PRT2527 with copanlisib/venetoclax in SSK41 and VL51 or **(C)** with ibrutinib in Karpas1718 and VL51 upon 72 hr of exposure to single agents or combinations, followed by MTT assay.

### The CDK9 gene is expressed in lymphoma samples

To further emphasize the importance of CDK9 as a therapeutic target for MZL lymphoma patients, we analyzed previously published datasets ^44-46^ and observed that CDK9 is expressed at levels comparable to those of pan-B cell markers, such as CD19 and CD21, across multiple cohorts of splenic MZL patients (Figure 6). We also confirmed CDK9 expression in FL, nodal, and extranodal MZL patient samples (Supplementary Figure S3). Lastly, in spleen samples from both non-tumoral individuals and splenic MZL patients, we found that CDK9 methylation is lower than in normal spleen tissue (Supplementary Figure S4), supporting the idea that reduced methylation contributes to CDK9 overexpression in tumor samples.

**Figure 6.**
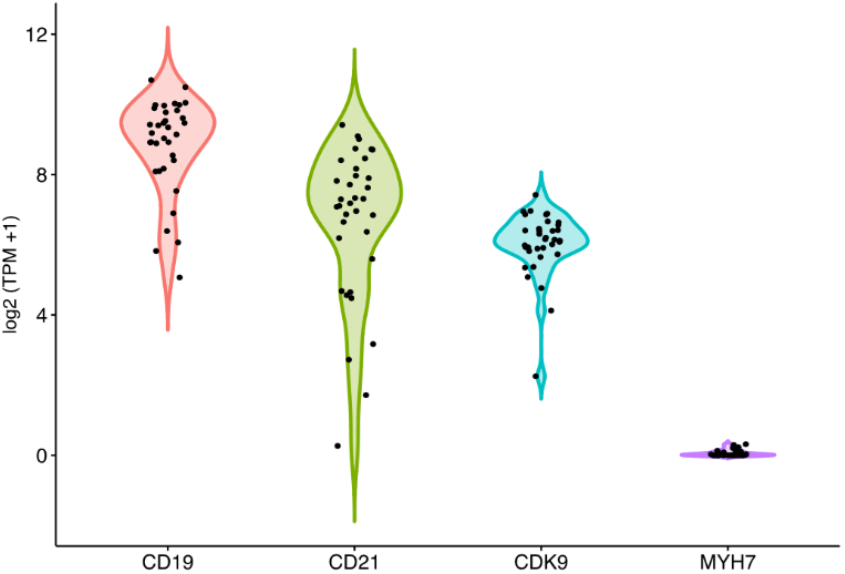
Violin plots showing the gene expression of *CDK9* in a cohort of splenic MZL patients (N=35), along with genes expected to be expressed (*CD19, CD21*) and not expressed (*MYH7*) in human spleen (dataset of clinical samples of splenic MZL patients ^44^).

## Discussion

Here, we demonstrated that i) the selective CDK9 inhibitor PRT2527 exhibits antitumor activity in MZL models, including cells with acquired resistance to BTK, PI3K, and BCL2 inhibitors, as a single agent and in combination, and ii) that it downregulates key oncogenic and survival proteins and increases expression of HLA class II molecules.

CDK9 is overexpressed in many cancers, making it a promising therapeutic target ^1-4^. In lymphomas, its expression has been reported, particularly in aggressive lymphomas ^4^. Although CDK9 expression has not been previously reported in MZL clinical specimens ^4^, we report here its expression at the RNA level across multiple datasets ^44-46^, suggesting that CDK9 could be a reasonable therapeutic target in this disease.

Inhibition of CDK9 with PRT2527 showed nanomolar cytotoxicity in nearly all tested parental and resistant cell lines. Its activity was consistent with that reported in diffuse large B-cell lymphoma models using PRT2527 or AZD5576, another CDK9 inhibitor ^13,31^. Notably, the effect was robust and, importantly, persisted in cell lines with secondary resistance to BTK, BCL2, and PI3K inhibitors. This suggests that CDK9 dependency is independent of the signaling pathways targeted by BTK, PI3K, or BCL2 inhibition. This aligns with previous findings in DLBCL and MCL, where CDK9 inhibition overcame resistance linked to changes in BCR signaling or apoptotic priming ^47^. The rapid loss of MCL1 following PRT2527 treatment is significant, as MCL1 is a key factor in resistance to venetoclax and BCR pathway inhibitors, and its depletion may explain the observed benefit of combination therapy.

Combination studies showed that PRT2527 enhanced the activity of BTK, PI3K, and BCL2 inhibitors. The strongest effects were observed with copanlisib plus venetoclax and ibrutinib, consistent with results reported with other CDK9 inhibitors ^16,47-49^. The synergy between PRT2527 and either copanlisib plus venetoclax or ibrutinib underscores the rationale for CDK9-based combinations that target survival signals and transcriptional addiction simultaneously. PI3K inhibitors lower prosurvival signaling and glucose metabolism, while BCL2 inhibition directly disrupts mitochondrial integrity. When combined with PRT2527’s rapid suppression of MCL1 and MYC, multiple apoptotic checkpoints are bypassed. Likewise, BTK inhibition reduces BCR-driven NF-κB signaling, and when paired with transcriptional repression, it further weakens the survival capacity of malignant B cells. Early results from clinical trials combining CDK9 inhibitors with BTK inhibitors (such as PRT2527/zanubrutinib ^28^ and tambiciclib/zanubrutinib (https://www.globenewswire.com/news-release/2025/02/20/3029678/0/en/SELLAS-Announces-Positive-Data-from-Phase-2a-Trial-of-SLS009-in-Combination-with-Zanubrutinib-in-DLBCL.html)) or BCL2 inhibitors (like tambiciclib/venetoclax ^26^, voruciclib/venetoclax ^24^, QHRD107/venetoclax/azacitidine ^27^) suggest that these combinations may be effective in MZL patients as well.

The transcriptomic data provided additional mechanistic insights. Downregulation of MYC, IRF4, and NF-κB–associated gene sets confirmed the expected on-target effects of CDK9 inhibition on short-lived transcripts. Induction of metabolic and oxidative phosphorylation pathways may reflect a stress response and possibly contribute to cell death, as recently reported in MCL exposed to the CDK9 inhibitor YX0798 ^14^. Notably, PRT2527 increased transcription of HLA class II genes and enhanced their surface expression, potentially improving the immune response of CD4^+^ and CD8^+^ T cells, as reported with HDAC inhibitors ^50^. The concurrent reduction of HLA class I could have complex immunological implications, possibly affecting cytotoxic T cell recognition while favoring NK cell activation ^51,52^. These immunomodulatory effects will require further investigation, especially in the context of immunotherapy combinations.

At the protein level, we confirmed a rapid and significant loss of MCL1 and MYC, aligning with the RNA-seq results and the short half-lives of these proteins, as previously reported ^12,13,15,16^. The observed suppression of RNA polymerase II and IRF4 further supports a broad impact on transcriptional programs essential for MZL survival. Interestingly, IRF4 downregulation occurred at low PRT2527 concentrations and did not increase at higher doses, suggesting a threshold-dependent effect or compensatory regulation.

In conclusion, our findings highlight the therapeutic potential of CDK9 inhibition in MZL, both as a single agent and in combination with other agents. PRT2527’s ability to overcome resistance to BTK, PI3K, and BCL2 inhibitors positions it as a promising option for salvage treatment in relapsed or refractory cases. Its dual influence on tumor-intrinsic survival pathways and immune gene expression creates opportunities to integrate it with targeted agents and immunotherapies. Given PRT2527’s favorable clinical tolerability profile, these preclinical results support further development of CDK9-based combinations for MZL and other B-cell malignancies.

## Acknowledgments

The authors used OpenAI’s ChatGPT (GPT-5, OpenAI, San Francisco, CA, USA) to assist in text preparation and drafting. All content was reviewed, edited, and verified by the authors, who take full responsibility for the manuscript.

## Potential Competing Interests

Alberto J. Arribas: travel grant from AstraZeneca and Floratek Pharma, advisory board fee from PentixaPharm.

Luciano Cascione: institutional research funds from Orion; travel grant from HTG.

Davide Rossi: grant support from Gilead, AbbVie, Janssen; honoraria from Gilead, AbbVie Janssen, Roche; scientific advisory board fees from Gilead, AbbVie, Janssen, AstraZeneca, MSD.

Anastasios Stathis: institutional research funds from Pfizer, MSD; Roche, Novartis, Amgen, Abbvie, Bayer, ADC Therapeutics, MEI Therapeutics, Philogen, Celestia. Astra Zeneca; travel grant from AbbVie and PharmaMar; consulting fee paid to institution from Jansen, Roche, Eli Lilly.

Diane Heiser: Prelude employment

Francesco Bertoni: institutional research funds from ADC Therapeutics, Bayer AG, BeiGene, Floratek Pharma, Helsinn, HTG Molecular Diagnostics, Ideogen AG, Idorsia Pharmaceuticals Ltd., Immagene, ImmunoGen, iOnctura, Mabtree, Menarini Ricerche, Nordic Nanovector ASA, Oncternal Therapeutics, Spexis AG; consultancy fee from BIMINI Biotech, Floratek Pharma, Helsinn, Immagene, Menarini, Vrise Therapeutics; advisory board fees to institution from Novartis; travel grants from Amgen, Astra Zeneca, iOnctura.

The other Authors have nothing to disclose.

## Authors’ contributions

EC, AJA: performed experiments, analyzed and interpreted data, performed data mining, prepared the figures, and co-wrote the manuscript.

LC performed data mining.

AR performed transcriptome profiling.

AS, DR: provided resources and advice.

DH: provided resources.

FB: designed research, interpreted data, and co-wrote the manuscript. All authors reviewed and accepted the final version of the manuscript.

