## Supplementary materials and methods for "CDK9 pharmacological inhibition with PRT2527 has antitumor activity in marginal zone lymphoma models and can improve the effects of BTK, PI3K, and BCL2 inhibitors"

### Supplementary tables and figures

**Supplementary Table S1. Drug concentration range and dilutions used in the single and combination experiments with the CDK9 inhibitor PRT2527. A)** 4-hour exposure to PRT2527, followed by washout and 48 hours of exposure to the second drug. MTT assay at the end of the treatment. **B)** 72 hours of exposure to single agents or combinations followed by MTT assay.

**A)**

| Experiment | Compound | cell line | max concentration | drug dilution |
|---|---|---|---|---|
| single | PRT2527 | all models | 10µM | 1/3 |
| combo with PRT2527 (0-10µM dil 1/3) | Copanlisib plus Venetoclax | SSK41, VL51 | Copanlisib 1µM<br>Venetoclax 10µM | 1/10 |
| combo with PRT2527 (0-10µM dil 1/3) | Idelalisib | Karpas1718, VL51 | 25µM | 1/5 |
| combo with PRT2527 (0-10µM dil 1/3) | Ibrutinib | VL51 | 25µM | 1/5 |

**B)**

| Experiment | Compound | cell line | max concentration | drug dilution |
|---|---|---|---|---|
| single | PRT2527 | SSK41, VL51, Karpas1718 | 1µM | 1/10 |
| combo with PRT2527 (0-1µM dil 1/10) | Copanlisib plus Venetoclax | SSK41, VL51 | Copanlisib 1µM<br>Venetoclax 10µM | 1/10 |
| combo with PRT2527 (0-1µM dil 1/10) | Ibrutinib | VL51, Karpas1718 | 10µM | 1/10 |

**Supplementary Table S2. Experimental conditions used for the RNA sequencing.** VL51 were exposed to the CDK9 inhibitor PRT2527 (IC50) or to DMSO for 4 hours. Then, RNA was extracted and processed for RNA sequencing.

| Experiment | Compound | cell line | Cell concentration | Drug concentration (IC50) |
|---|---|---|---|---|
| RNA-Seq | PRT2527 | VL51 | 500,000/well | 850nM |

**Supplementary Table S3. Transcriptomic data included as a separate excel file.**

**Supplementary Figure 1. Surface HLA expression in VL51 parental cells after exposure to PRT2527.** Cells were exposed for 24 hours to increasing concentrations of PRT2527 (100 nM, 400 nM, or 850 nM) or DMSO (control). Ridgeline plot representative of at least two independent experiments.

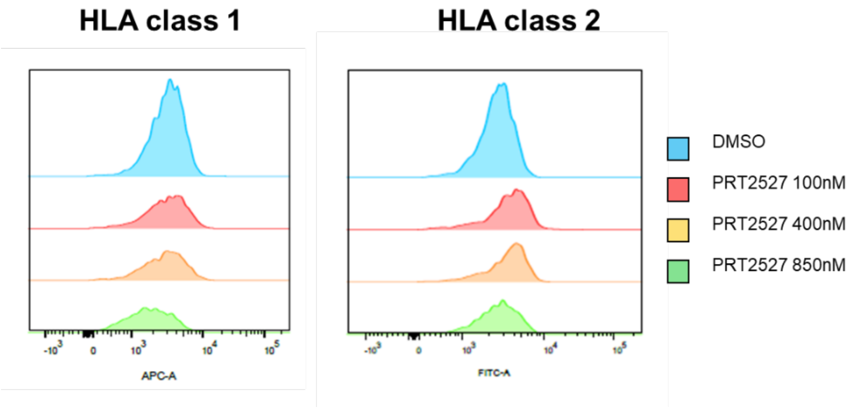


**Supplementary Figure 2. Surface HLA expression in HAIR-M, ESKOL, and Karpas1718 cell lines after exposure to PRT2527.** Cells were exposed for 24 hours to a concentration of PRT2527 corresponding to the IC50 (310 nM for HAIR-M, 185 nM for ESKOL, or 850 nM for Karpas1718) or DMSO (control). **A)** Bar plots represent the mean of at least two independent experiments, error bars for standard deviation. **B)** Histograms representative of at least two independent experiments.

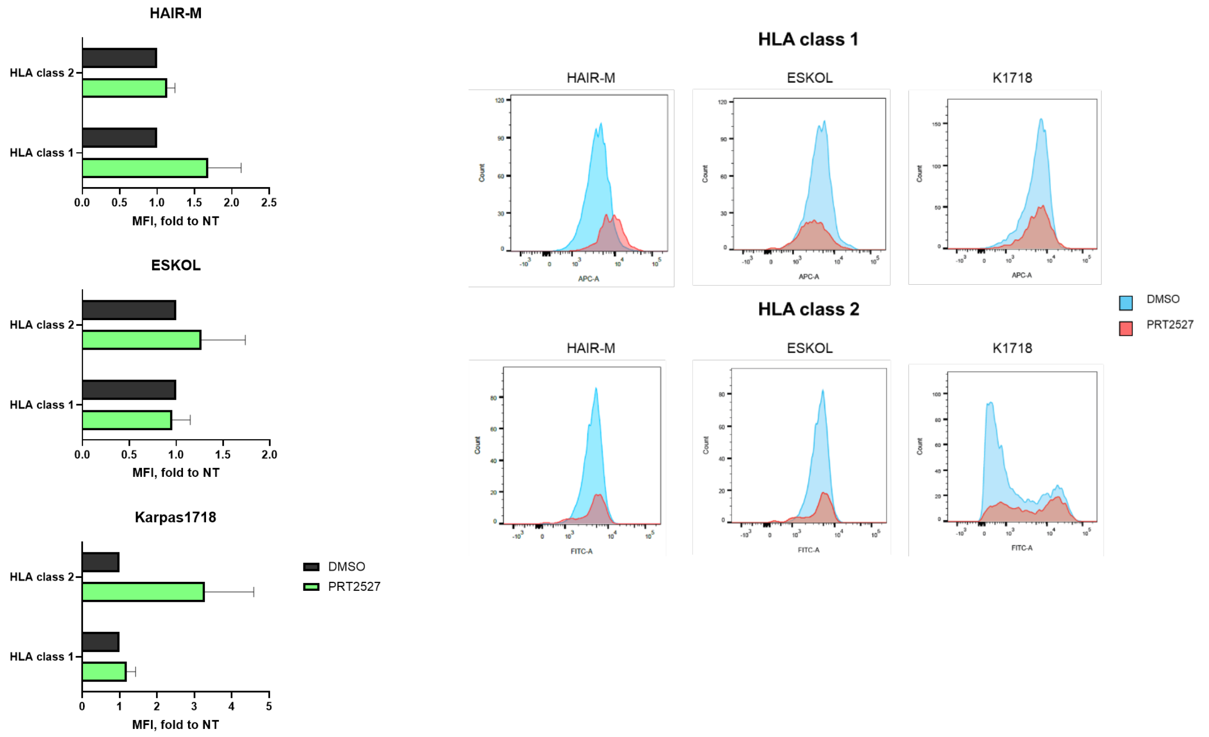

**Supplementary Figure 3. Boxplots showing the gene expression of *CDK9* in a cohort of patients, along with genes expected to be expressed (*CD19*) and not expressed (*MYH7*) in lymphoma.** Dataset of clinical samples [1] of FL (n = 16), extranodal MZL (MALT lymphomas) (= 5), nodal MZL (NMZL, n = 15), and splenic MZL (SMZL, n = 4) patients compared to reactive lymph nodes (RLN, n = 8).

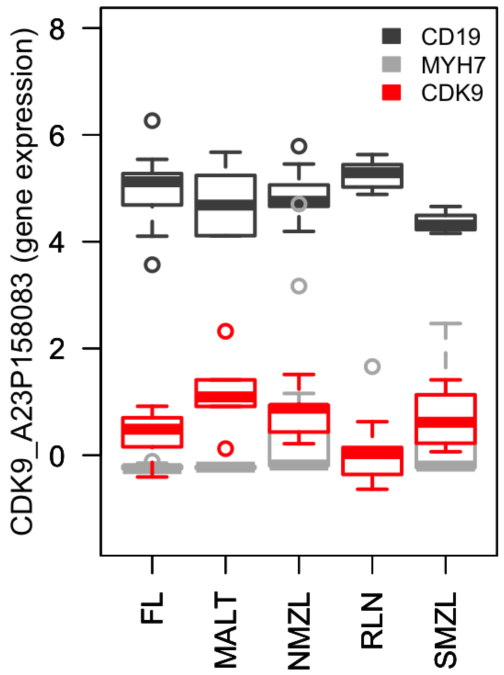

**Supplementary Figure S4. Boxplot showing the B-value of the *CDK9* methylation. Dataset of clinical samples [2] of splenic MZL (SMZL) patients (n = 126) and non-tumoral samples from the spleen (n = 3).**

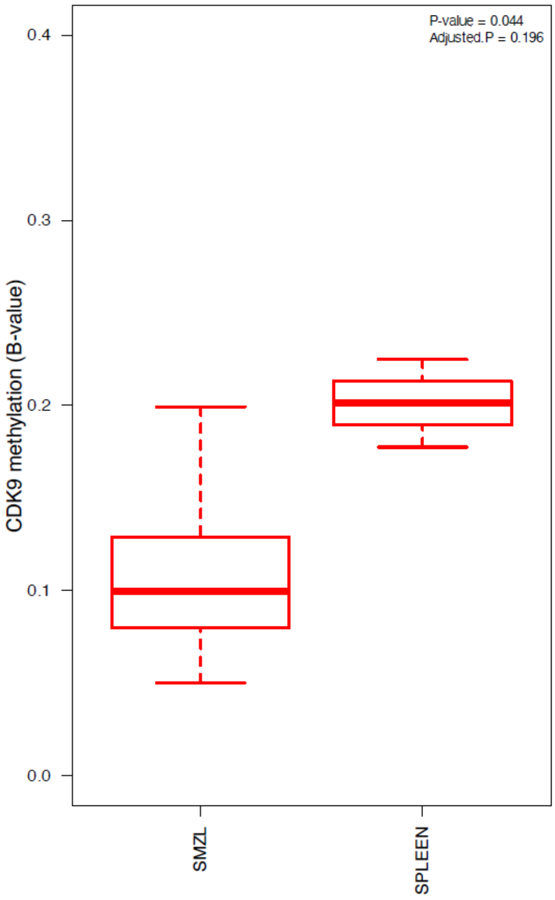
